# Barrier effects on the spatial distribution of *Xylella fastidiosa* in Alicante, Spain

**DOI:** 10.1101/2021.04.01.438042

**Authors:** Martina Cendoya, Ana Hubel, David Conesa, Antonio Vicent

## Abstract

Spatial models often assume isotropy and stationarity, implying that spatial dependence is direction invariant and uniform throughout the study area. However, these assumptions are violated when dispersal barriers are present in the form of geographical features or disease control interventions. Despite this, the issue of non-stationarity has been little explored in the context of plant health. The objective of this study was to evaluate the influence of different barriers in the distribution of the quarantine plant pathogenic bacterium *Xylella fastidiosa* in the demarcated area in Alicante, Spain. Occurrence data from the official surveys in 2018 were analyzed with four spatial Bayesian hierarchical models: i) a stationary model representing a scenario without any control interventions or geographical features; ii) a model with mountains as physical barriers; iii) a model with a continuous or iv) discontinuous perimeter barrier as control interventions surrounding the infested area. Barriers were assumed to be totally impermeable, so they should be interpreted as areas without host plants and in which it is not possible for infected vectors or propagating plant material to pass through. Inference and prediction were performed through the integrated nested Laplace approximation methodology and the stochastic partial differential equation approach. In the stationary model the posterior mean of the spatial range was 4,030.17 m 95% CI (2,907.41, 5,563.88), meaning that host plants that are closer to an infected plant than this distance would be at risk for *X. fastidiosa*. This distance can be used to define the buffer zone around the infested area in Alicante. In the non-stationary models, the posterior mean of the spatial range varied from 3,860.88 m 95% CI (2,918.61, 5,212.18) in the mountain barrier model to 6,141.08 m 95% CI (4,296.32, 9,042.99) in the continuous barrier model. Compared with the stationary model, the perimeter barrier models decreased the probability of *X. fastidiosa* presence in the area outside the barrier. Differences between the discontinuous and continuous barrier models showed that breaks in areas with low sampling intensity resulted in a higher probability of *X. fastidiosa* presence. These results may help authorities prioritize the areas for surveillance and implementation of control measures.

## Introduction

The plant pathogen *Xylella fastidiosa* is a xylem-limited bacterium with a host range of more than 500 plant species (EFSA 2020). Six subspecies and a number of sequence types (STs) have been described in *X. fastidiosa*, with different genetic traits, host ranges and aggressiveness (Denancé et al. 2017). This pathogen was confined to the American continent for decades (Janse and Obradovic 2010), but in 2013 it was first reported in Europe associated with a disease causing serious losses in olives in southern Italy (Schneider et al. 2020). Since then, *X. fastidiosa* has been detected in France, Spain, Portugal and Israel, affecting multiple host plants in agricultural and natural settings. To date, the subspecies *pauca*, *fastidiosa* and *multiplex* have been reported in the Mediterranean Basin (EFSA 2019). *X. fastidiosa* is regulated in the European Union (EU) as a quarantine pest (i.e. pathogen) under Regulation (EU) 2016/2031 and Commission Implementing Regulation (EU) 2019/2072. It is also included in the list of priority pests for the EU by Commission Delegated Regulation (EU) 2019/1702.

Two groups of xylem sap-feeding insects have been identified as the natural means by which *X. fastidiosa* spreads: sharpshooters (Cicadellidae family, Cicadellinae subfamily) and spittlebugs (Aphrophoridae, Cercopidae and Clastopteridae families) (Almeida et al. 2005; Almeida and Nunney 2015). Nymphs and adults of the vectors acquire the bacteria by feeding on the xylem of infected plants. Bacteria then multiply in the insect foregut, but vectors lose infectivity with molting. Adult vectors can inoculate healthy plants immediately after acquisition and throughout their whole lifetime, although the bacterium is not transmitted to the progeny (Almeida and Purcell 2006).

The pathogen can be introduced and further spread into new areas with infected plant material for planting or grafting (EFSA 2015). Genetic studies indicated that the *X. fastidiosa* subsp. *pauca* ST53 strain CoDiRO, which is decimating olive trees in Italy, originated in Central America (Giampetruzzi et al. 2017) and was probably introduced with infected coffee plants imported as ornamentals. Phylogenetic analyses also indicated that *X. fastidiosa* subsp. *multiplex* ST81 and *X. fastidiosa* subsp. *fastidiosa* ST1 were introduced into the island of Majorca, Spain, with infected almond graftings from California, US (Moralejo et al. 2020). Similarly, *X. fastidiosa* subsp. *multiplex* ST6 in Alicante, Spain, and Corsica, France, as well as ST87 from Tuscany, Italy, might have been introduced from California (Landa et al. 2020). Human-assisted movement of infected insect vectors on plants or on their own as ‘hitch-hikers’ in vehicles can also disseminate *X. fastidiosa*, though information on these means of spread is limited (EFSA 2015).

Species distribution models (SDMs) are widely used to associate the geographic settings of species with biotic and abiotic factors, establish favorable areas for the expansion of populations, develop risk maps for the potential establishment of pathogens, and predict the distribution of species in space and time, among others (Martínez-Minaya et al. 2018). These types of models can be developed with different methodologies, such as generalized linear models (GLM), generalized additive models (GAM), neural networks, maximum entropy models (e.g. Maxent) and climate envelope models (e.g. Bioclim). The literature available on the applications and methodologies of SDMs is quite extensive, with some reviews such as Guisan and Zimmermann (2000), Elith and Leathwick (2009) or Martínez-Minaya et al. (2018) that compile and describe the different modeling approaches.

Several studies have been conducted on the potential distribution of *X. fastidiosa* associated with climatic factors (Bosso et al. 2016; Godefroid et al. 2019; Hernández and García 2019; EFSA 2019). However, most of these models are based on the assumption that observations are independent, without taking into account the spatial dependence that often exists among the geographical locations. Failing to consider spatial correlation may lead to an overestimation of the model parameters and thus inaccurate results (Latimer et al. 2006). Advances in computational methods have made it possible to implement more complex models and, hence, a more straightforward incorporation of spatial dependencies in SDMs (Blangiardo and Cameletti 2015). Among these advances, here we will focus on hierarchical Bayesian models, which allow random effects and complex dependency structures to be incorporated easily taking into account all the non-observed uncertainties (Banerjee et al. 2004; Blangiardo and Cameletti 2015).

An additional and often overlooked problem in the analysis of spatial data is that models usually assume stationarity (i.e., the spatial effect is invariant to the map translation) and isotropy (i.e., the spatial effect is invariant to the map rotation), that is, the autocorrelation between two locations only depends on the Euclidean distance. However, relying on these two assumptions can produce misleading results, with unrealistic associations and/or bias in the prediction of the species distribution, when elements such as barriers that are an obstacle to the movement of the species are present in the study area. To address this issue, Bakka et al. (2019) introduced an approach that makes it possible to deal with non-stationary spatial processes where, as in our study, stationary also includes isotropy for convenience. This approach has been applied in marine species distribution studies, where the coastline was implemented as a physical barrier. In particular, in the above-mentioned work Bakka et al. (2019) modeled the distribution of fish larvae in the Finnish Archipelago (Finland), while Martínez-Minaya et al. (2019) conducted a study on the seasonal distribution of bottlenose dolphins in the Archipelago de La Maddalena (Italy).

Barriers are an intrinsic part of the principles of plant disease control, i.e. exclusion, eradication, protection and resistance (Maloy 1993). Exclusion strategies aim to prevent the pathogens from entering new areas. Barriers in the form of prohibitions restricting the import of plants, interceptions through border inspections and subsequent elimination of the pathogen are enforced by legal provisions worldwide. In the case of *X. fastidiosa*, the Commission Implementing Regulation (EU) 2020/1201 establishes special requirements for the import of host plants from third countries into the EU. When exclusion fails, eradication is attempted by removing the infected plants to limit further spread of the disease. According to Commission Implementing Regulation (EU) 2020/1201, demarcated areas consisting of an infected (i.e. infested) zone and a buffer zone should be established for *X. fastidiosa*. Eradication measures should then be implemented to ensure the removal of the infected plants and control of vector populations. Special requirements are also set for the movement of specified plants from the demarcated area.

Protection from already established diseases can be accomplished with barriers such as screenhouses, plastic covers and distance from inoculum sources that prevent pathogens and vectors from contacting host plants. Windbreaks can also prevent the movement of pathogen propagules. In areas where *X. fastidiosa* is endemic, screen and planting barriers have been evaluated to reduce vector spread (Daugherty and Almeida 2009; Blua et al. 2005). Finally, plant resistance limits the infection and multiplication of plant pathogens, acting as a barrier for the onset of disease epidemics. In this regard, recent advances have been made to obtain grapevine and olive cultivars that are resistant to *X. fastidiosa* (Krivanek et al. 2006; Giampetruzzi et al. 2016).

All these examples described above illustrate to what extent the presence of barriers and their resulting non-stationarity can shape the spatial dimension of plant disease epidemics. Nevertheless, apart from performing separate directional spatial autocorrelation analyses to study whether a process is isotropic (Madden et al. 2007), the issue of non-stationarity has been scarcely explored in the context of plant disease epidemiology.

Our study focuses on the demarcated area for *X. fastidiosa* in Alicante, Spain. The pathogen was first reported in this region in 2017 and since then has been under official control in accordance with EU legislation. In Alicante, *X. fastidiosa* subsp. *multiplex* ST6 was identified as affecting mainly almond trees (*Prunus dulcis*). The two insect species where *X. fastidiosa* has been detected in this area are *Philaeuns spumarius* L. (Hemiptera: Aphrophoridae) and *Neophilaenus campestris* Fallen (Hemiptera: Aphrophoridae) (GVA 2020). Despite its relatively small extension, the study area of Alicante presents a great orographic diversity, from the sea level to mountain ranges rising to an altitude of above 1,500 m. This particular geographic setting must be taken into account to model the occurrence of *X. fastidiosa*, since it can determine the presence of host plants and also affect the behavior of the vectors, thus violating the stationarity and isotropy assumption. In addition to these geographic barriers, the control measures for *X. fastidiosa* established by the EU legislation are aimed at limiting the spread of the disease, which also represents a potential dispersal barrier to be considered.

With all this in mind, the aim of this study is to describe how the presence of different kinds of barriers produce different results in terms of predicting the presence of a species. In particular, the occurrence of *X. fastidiosa* in Alicante was analyzed with four spatial modeling scenarios, three of them including dispersal barriers.

## Methods

### Database

The georeferenced data from the official surveys carried out for *X. fastidiosa* in Alicante in 2018 were provided by the plant health authority (*Sanitat Vegetal, Generalitat Valenciana*). This database contained the plant species sampled, the result of the laboratory analysis being positive (i.e. presence) or negative (i.e. absence) for *X. fastidiosa* based on real-time PCR (EPPO 2019), as well as the UTM coordinates of the location where the sample was taken.

Samples were also collected from plant species that were not known to be natural hosts for the *X. fastidiosa* subsp. *multiplex* strains present in the study area, such as *Olea europaea*, of which 2,414 samples were collected during that period, all of them resulting negative for *X. fastidiosa*. In order to avoid biases in the estimation due to this large number of negative samples from non-host species, only the samples from plant species having at least one positive for *X. fastidiosa* were considered for further analysis. The plant species selected were: *Prunus dulcis*, *P. armeniaca*, *P. domestica*, *Calicotome spinosa*, *Rhamnus alaternus*, *Phagnalon saxatile*, *Helichrysum italicum*, *Polygala myrtifolia*, *Rosmarinus officinalis* and *Laurus nobilis*. The dataset consisted of a total of 4,205 samples, 1,151 were positive and 3,054 were negative for *X. fastidiosa*, distributed in the demarcated area of Alicante with an extension of approximately 1,346 km^2^ (GVA 2019) (Fig. S1).

### Geostatistical model

Considering the georeferenced data as observations made at continuous locations occurring within a defined spatial domain, they were classified as geostatistical data. One of the characteristics of this type of spatial data is that the main objective of its analysis is to enable prediction within the study region (Cressie 1993). A point-referenced spatial hierarchical model (Diggle et al. 1998) was used to model the geostatistical data, while inference and prediction were performed within the Bayesian paradigm. As posterior distributions of the parameters and hyperparameters, along with the posterior predictive distributions of the predicted values in unobserved locations, do not have analytical expressions, the integrated nested Laplace approximation (INLA) methodology (Rue et al. 2009) was used to numerically approximate them.

Defining a hierarchical Bayesian spatial model can be seen as a three-step process. Firstly, a probability distribution must be identified for the observations available at the spatial locations. In this case, it was assumed that *y_i_*, the occurrence of *X. fastidiosa* at location *i*, follows a Bernoulli distribution (1 indicating presence and 0 absence), that is, *y_i_ ∼* Bernoulli(*π_i_*), where *π_i_* represents the probability of presence at location *i*. In a second step, this probability of presence *π_i_* is linked (usually via the logit link when the response is Bernoulli) to a linear predictor and a latent Gaussian random field, whose covariance matrix Σ depends on two hyperparameters: the variance *σ*^2^ and the range *r* of the spatial effect. Finally, the third step consists in assigning the corresponding priors and hyperpriors of the parameters and hyperparameters of the model. Despite its wide acceptance, INLA cannot be directly applied when dealing with continuously indexed Gaussian fields (GF). The underlying reason is that the cost of factorizing dense covariance matrices can be computationally demanding. Lindgren et al. (2011) proposed an alternative approach by using an approximate stochastic weak solution to a Stochastic Partial Differential Equation (SPDE) as a Gaussian Markov random field (GMRF) approximation to a continuous GF with Matérn covariance structure. A GMRF is a discretely indexed GF characterized by a sparse precision matrix *Q*, the factorizing computational cost of which is of order *O*(*n*^3^*^/^*^2^), a large computational improvement compared to the factorization of a dense covariance matrix (of order *O^n^*) that would imply the GF. In the approach proposed by Lindgren et al. (2011), the finite element method provides a solution to the SPDE, through the construction of a *mesh* (Appendix S1: Fig. S2a), which consists in the triangulation of the study area (Bakka et al. 2018).

Using this approximation, the spatial term is reparameterized as *u ∼ N* (0*, Q^−^*^1^(*κ, τ*)), where the parameters *κ* and *τ* control the range (*r*) and the variance 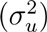. Specifically, 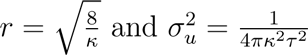 (Lindgren et al. 2011). However, for a more intuitive interpretation, the spatial effect was parameterized in terms of the marginal standard deviation and the range (Krainski et al. 2019).

Therefore, the hierarchical Bayesian spatial model with the Krainski et al. (2019) reparameterization can be expressed as:

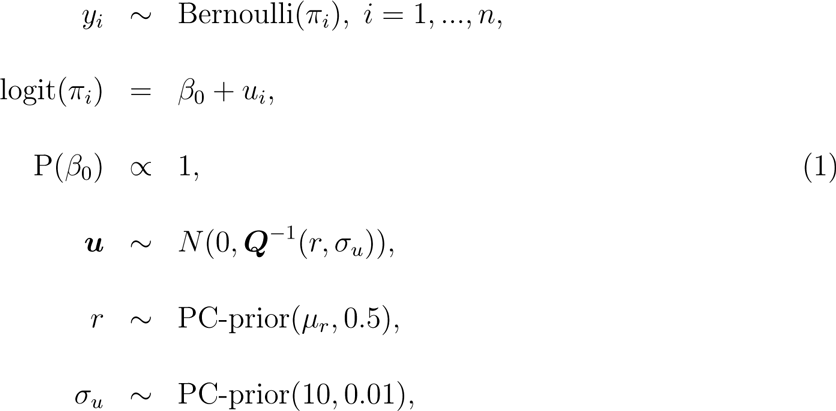

where *π_i_* is the probability of the presence of *X. fastidiosa* at location *i*, *β*_0_ is the intercept, and ***u*** is the spatial effect. As can be observed, the linear predictor was reduced just to the intercept, the underlying reason being that previous works had indicated a dominating effect of the spatial component compared to available covariates in the demarcated area (Cendoya et al. 2020). This model already includes the scarce prior knowledge about parameters, expressed via a non-informative improper prior for the intercept, and about the hyperparameters. In this latter case, following Fuglstad et al. (2019), Penalized Complexity priors (PC-priors) were used to express vague prior knowledge about them. In particular, a PC-prior for the range was defined as P(*r < µ_r_*) = 0.5, where *µ_r_* was chosen as 50% of the diameter of the study region, while a PC-prior P(*σ_u_ >* 10) = 0.01 was defined for the standard deviation of the spatial effect.

### Non-stationarity

The model introduced in the previous subsection assumes stationarity and isotropy. In order to deal with non-stationarity (i.e., non-stationary and anisotropic spatial processes), the approach presented by Bakka et al. (2019) was used. As happens in stationary models, estimating and predicting in non-stationary models can be rather complicated. In their proposal, Bakka et al. (2019) approximated them also by means of the SPDE approach using the finite element method. However, in this case a system of two SPDEs is presented, one for the barrier area and the other for the remaining area, which we have also denominated as the normal area, adapting their terminology.

In particular, a non-stationary spatial effect *u*(*s*) is the solution to the following system of stochastic differential equations:

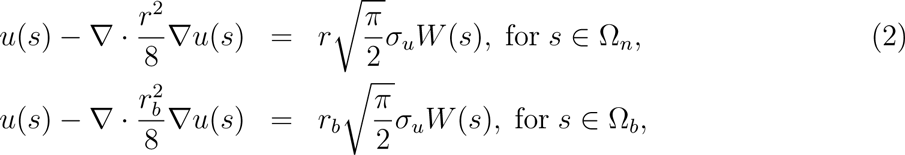

where *u*(*s*) is the spatial effect, Ω*_n_* is the normal area and Ω*_b_* is the barrier area. *r* and *r_b_* are the ranges for the normal and barrier areas, respectively. *σ_u_* is the marginal standard deviation, 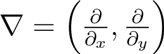 and *W* (*s*) denotes white noise. Note that in the barrier area the correlation is eliminated by introducing a different Matérn field, with the same standard deviation, but with a range close to zero.

### Models

In order to analyze the effect of including barriers on the occurrence of *X. fastidiosa* in the study area, the following models were performed and compared:

1. **Stationary model**. Model in which both stationarity and isotropy are assumed, without any barrier. This model represents a scenario without any disease control interventions or geographical features potentially affecting the spread of the pathogen (Fig. 1a).
2. **Mountain barrier model**. Non-stationary model with barriers defined by the areas over 1,065 m, the maximum altitude where a sample positive for *X. fastidiosa* was found in the study area. This model represents a scenario without any disease control interventions but with geographical features impeding the spread of the pathogen (Fig. 1b).
3. **Continuous barrier model**. Non-stationary model with a continuous barrier surrounding the infested area. This barrier consisted of a perimeter band 1,000 m wide, 500 m away from the outermost samples that were positive for *X. fastidiosa*. The width of the barrier was fixed to be lower than the range estimated for the stationary model. This model represents a cordon sanitaire where all host plants were removed and measures implemented to completely impede the spread of *X. fastidiosa*. For consistency, the perimeter band was also implemented along the coastline (Fig. 1c).
4. **Discontinuous barrier model**. The same non-stationary model described above but with a discontinuous barrier surrounding the infested area. In this case, breaks of different sizes (1,000-3,200 m) were made in the perimeter band, facing sampled and non-sampled areas outside the barrier. This model represents a cordon sanitaire where all host plants were removed, but measures to impede the spread of *X. fastidiosa* have been implemented only in some parts (Fig. 1d).

**Figure 1.**
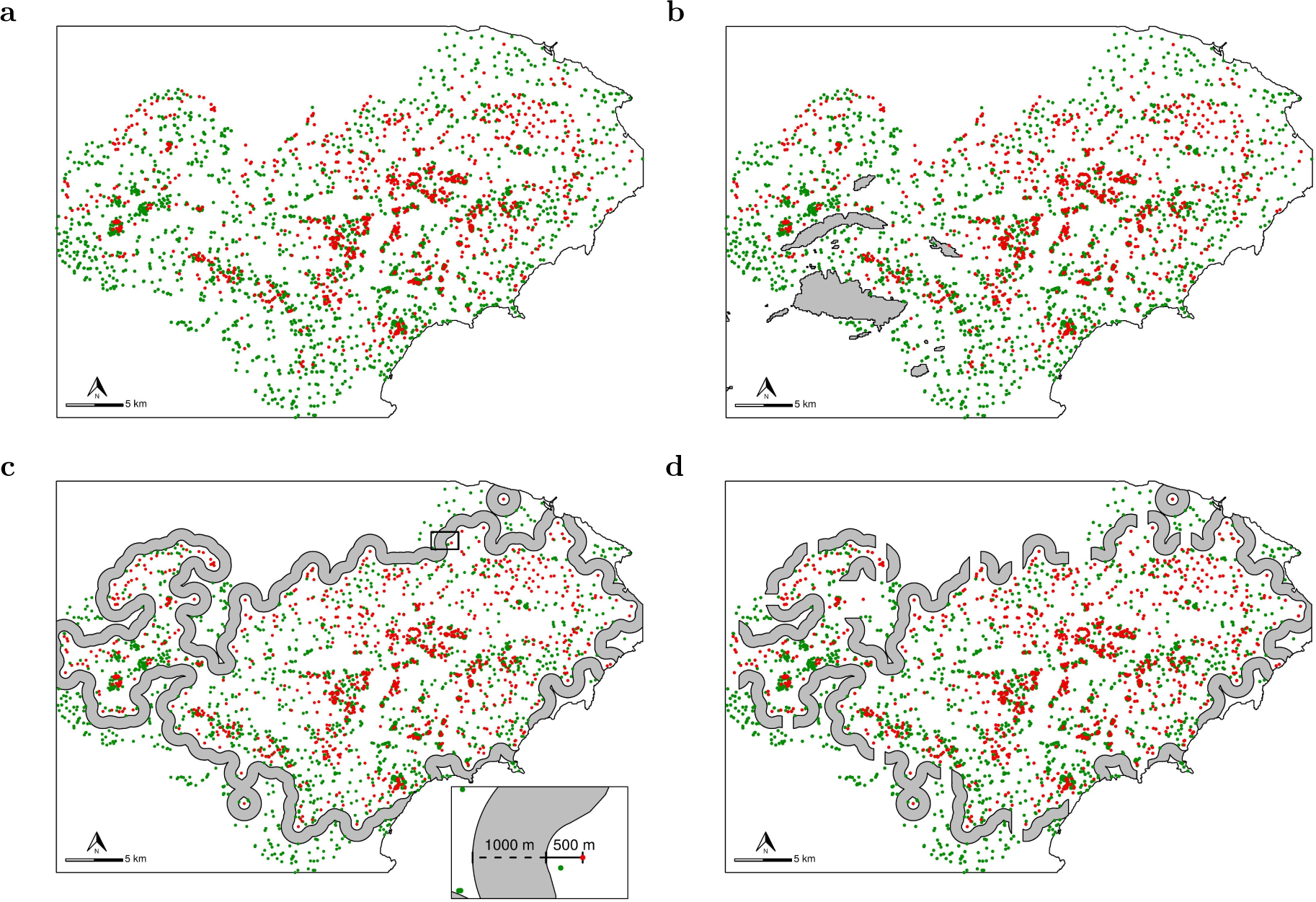
Positive (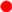) and negative (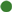) samples for *Xylella fastidiosa* and barriers incorporated into each model (shaded area). (a) Stationary model, without barriers; (b) mountain barrier model; (c) continuous barrier model; and (d) discontinuous barrier model.

In the non-stationary models (ii, iii and iv), following Eq. 2, Ω*_b_* represented the area occupied by the barriers, i.e., the area above 1,065 m in the mountain barrier model and the area of the cordon sanitaire in continuous and discontinuous barrier models. Ω*_n_* included the remaining area in each model.

All models were fitted using the INLA methodology with the R-INLA package (http://www.r-inla.org) for R software (R Core Team 2021). For each model a *mesh* was built, specifying in each case the barrier areas (Appendix S1: Fig. S2). In the three non-stationary models, observations in the barriers were eliminated (all of them negative samples), following the assumption that *X. fastidiosa* cannot be present in this specific area.

Differences between the stationary model and those with barriers, along with the differences between the discontinuous and continuous ones, were obtained by subtracting the means of their corresponding posterior predictive distributions.

## Results

In the stationary model, the posterior mean of the intercept was −1.95 in the linear predictor scale. Taking into account that when the spatial effect is zero, the mean posterior probability of the presence of *X. fastidiosa* is equivalent to the exponential transformation of the intercept, in this case, the probability of presence given only by the intercept was 0.14. The posterior mean of the spatial range was 4,030.17 m, with a 95% credible interval (CI) (2,907.41, 5,563.88) (Table 1). Therefore, we assume that two observations separated by more than this distance were not spatially correlated, that is, they are independent.

**Table 1:** Mean and 95% credible interval (CI) for the intercept (*β*_0_) and hyperparameters (*r* and *σ_u_*) of the models. *β*_0_ is the intercept, *r* is the range and *σ_u_* is the standard deviation of the spatial effect.

|  |  | Mean | 95% CI |
| --- | --- | --- | --- |
| Stationary | $\beta_0$ | -1.68 | (-2.21, -1.23) |
| | $r$ | 4030.17 | (2907.41, 5563.88) |
| | $\sigma_u$ | 1.52 | (1.28, 1.80) |
| Mountain<br>barrier | $\beta_0$ | -1.61 | (-2.09, -1.19) |
| | $r$ | 3860.88 | (2918.61, 5212.18) |
| | $\sigma_u$ | 1.43 | (1.20, 1.71) |
| Continuous<br>barrier | $\beta_0$ | -1.79 | (-2.63, -1.15) |
| | $r$ | 6141.08 | (4296.32, 9042.99) |
| | $\sigma_u$ | 1.50 | (1.20, 1.88) |
| Discontinuous<br>barrier | $\beta_0$ | -1.57 | (-2.23, -1.04) |
| | $r$ | 5298.90 | (3813.16, 7557.78) |
| | $\sigma_u$ | 1.44 | (1.17, 1.78) |

The posterior mean of the intercept in the mountain barrier, continuous barrier and discontinuous barrier models was −1.88, −2.07 and −1.86, respectively (Table 1). Therefore, in areas where there was no influence of the spatial effect, through the exponential transformation of these values, a probability of presence of the pathogen of 0.15 was obtained with the mountain barrier model, 0.13 with the continuous barrier model and 0.16 with the discontinuous barrier model.

In the mountain barrier model, a posterior mean of the spatial range of 3,860.88 m was obtained with a 95% CI (2,918.61, 5,212.18). In the continuous barrier model the range was greater than in the previous case and with more variability, obtaining a posterior mean of 6,141.08 m, with a 95% CI (4,296.32, 9,042.99). The estimation of the discontinuous barrier model resulted in a posterior mean of the range of 5,298.90 m with a 95% CI (3,813.16, 7,557.78) (Table 1).

The Matérn correlation function represents the spatial correlation between two observations as a function of distance, where the range is the distance from which two observations can be considered independent (Cressie 1993). The Mátern correlation function was estimated in each model using the posterior mean of the range obtained. The function was similar in the stationary and mountain barrier models, where the spatial correlation decreases quickly in the first 4,000 m. In the continuous barrier and discontinuous barrier models, the spatial correlation as a function of distance had a more gradual decrease due to the greater range obtained in the estimation (Fig. 2).

**Figure 2.**
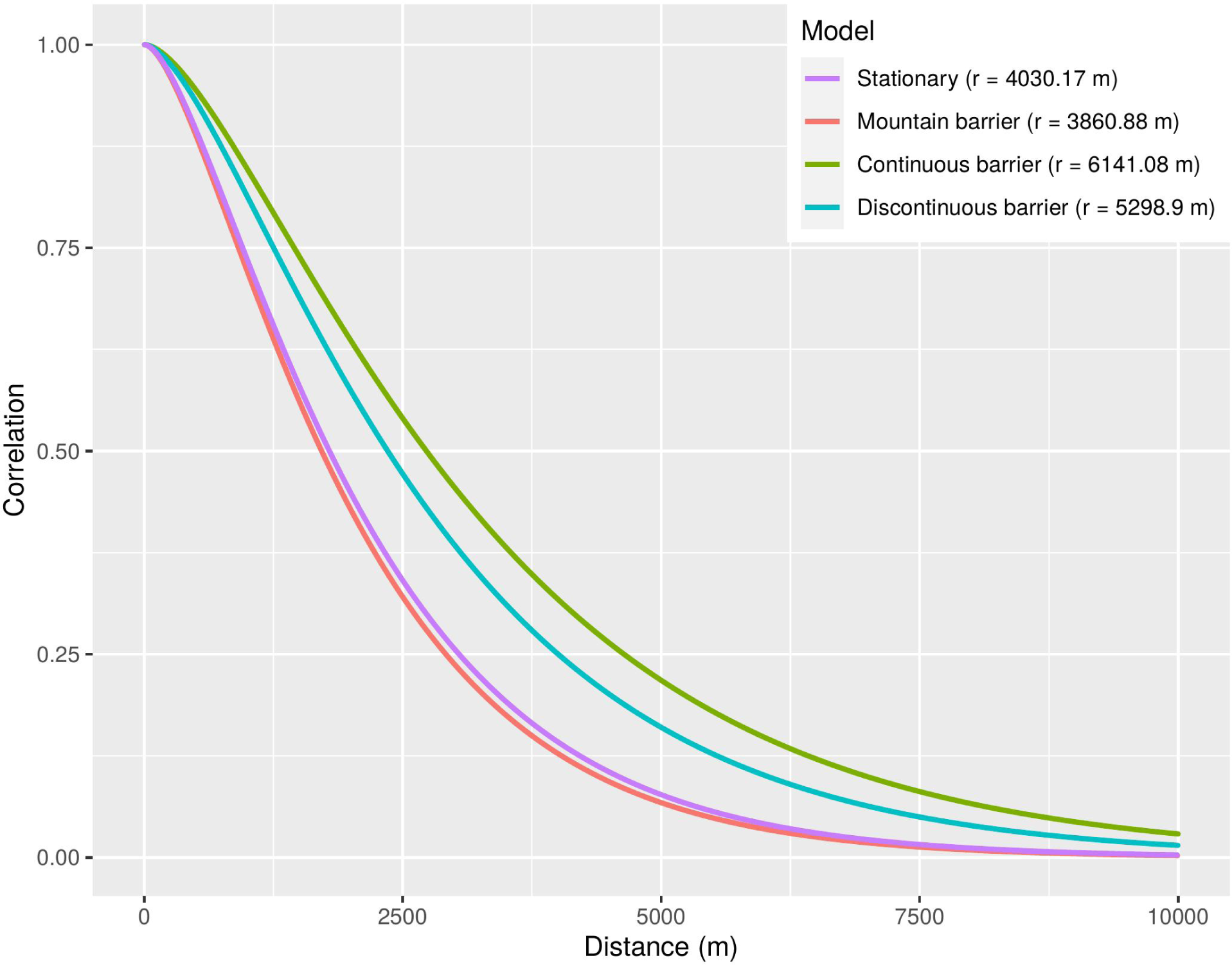
Representation of the Matérn correlation function for the posterior mean of the range obtained in each model.

Given the model described in Eq. 1, the mean of the posterior predictive distribution, expressed in terms of probability, was defined by the intercept and the spatial effect. In general, in the four modeling scenarios the probability of the presence of *X. fastidiosa* was higher in the areas where the positive samples were concentrated, being close to zero in the areas where negative samples predominated. In the non-sampled areas at distances from the observations outside the range, and thus without any influence of the spatial effect, the probability of the presence of *X. fastidiosa* only depended on the intercept. Regarding the standard deviation of the posterior predictive distribution, higher values were obtained in the non-sampled areas, while the sampled areas where *X. fastidiosa* was not detected showed very low variability (Fig. 3).

**Figure 3.**
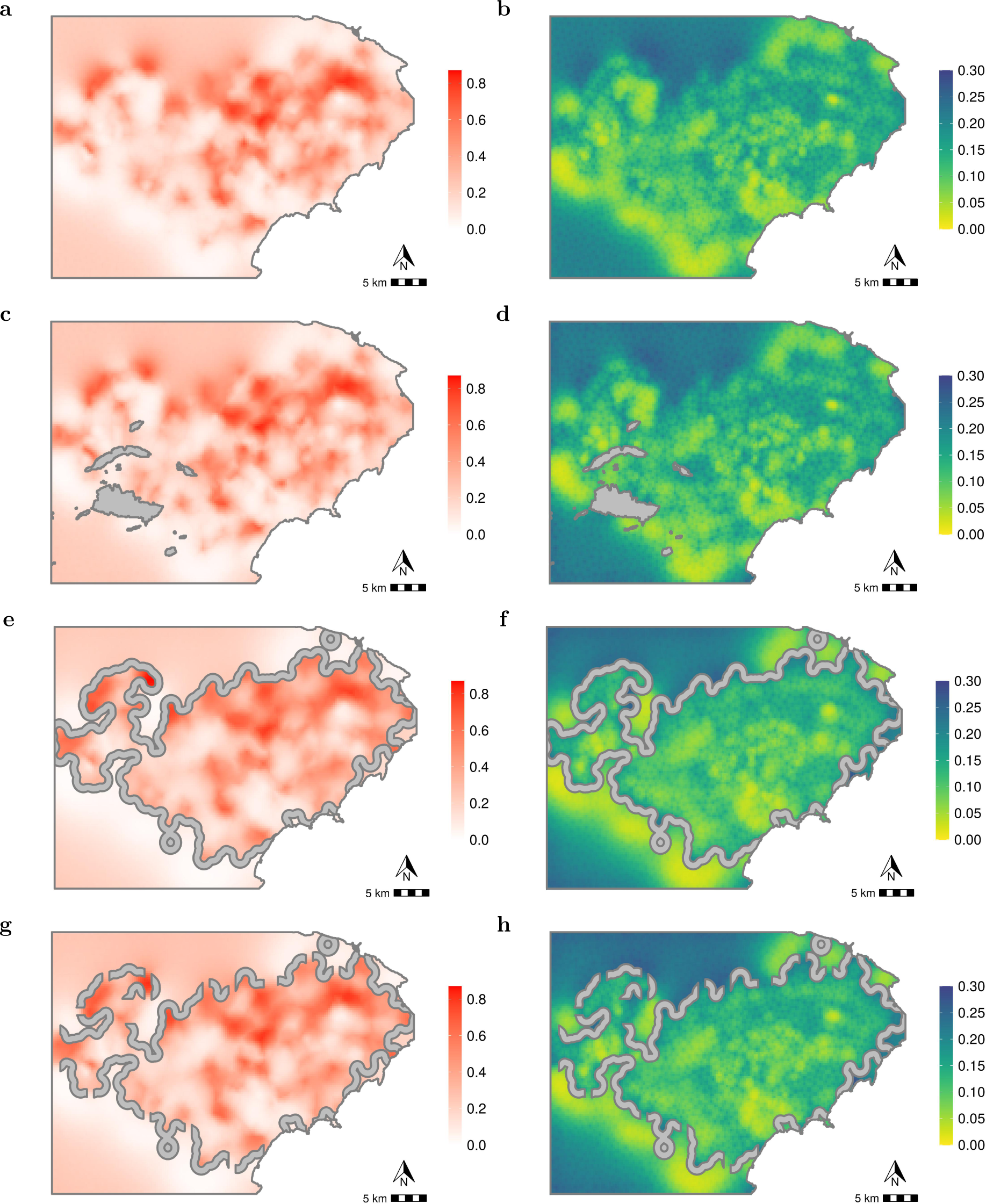
Mean (left) and standard deviation (right) of the posterior predictive distribution of the probability of *Xylella fastidiosa* presence for each model. (a, b) Stationary model; (c, d) mountain barrier model; (e, f) continuous barrier model; and (g, h) discontinuous barrier model.

The range of values of the mean and standard deviation of the posterior predictive distribution was similar in all four models (Fig. 3). However, the difference between the mean of the stationary model and the mountain barrier model was negative in the area around the barrier (Fig. 4a). This implies that the probability of *X. fastidiosa* presence in those areas was higher in the mountain barrier model than in the stationary model.

**Figure 4.**
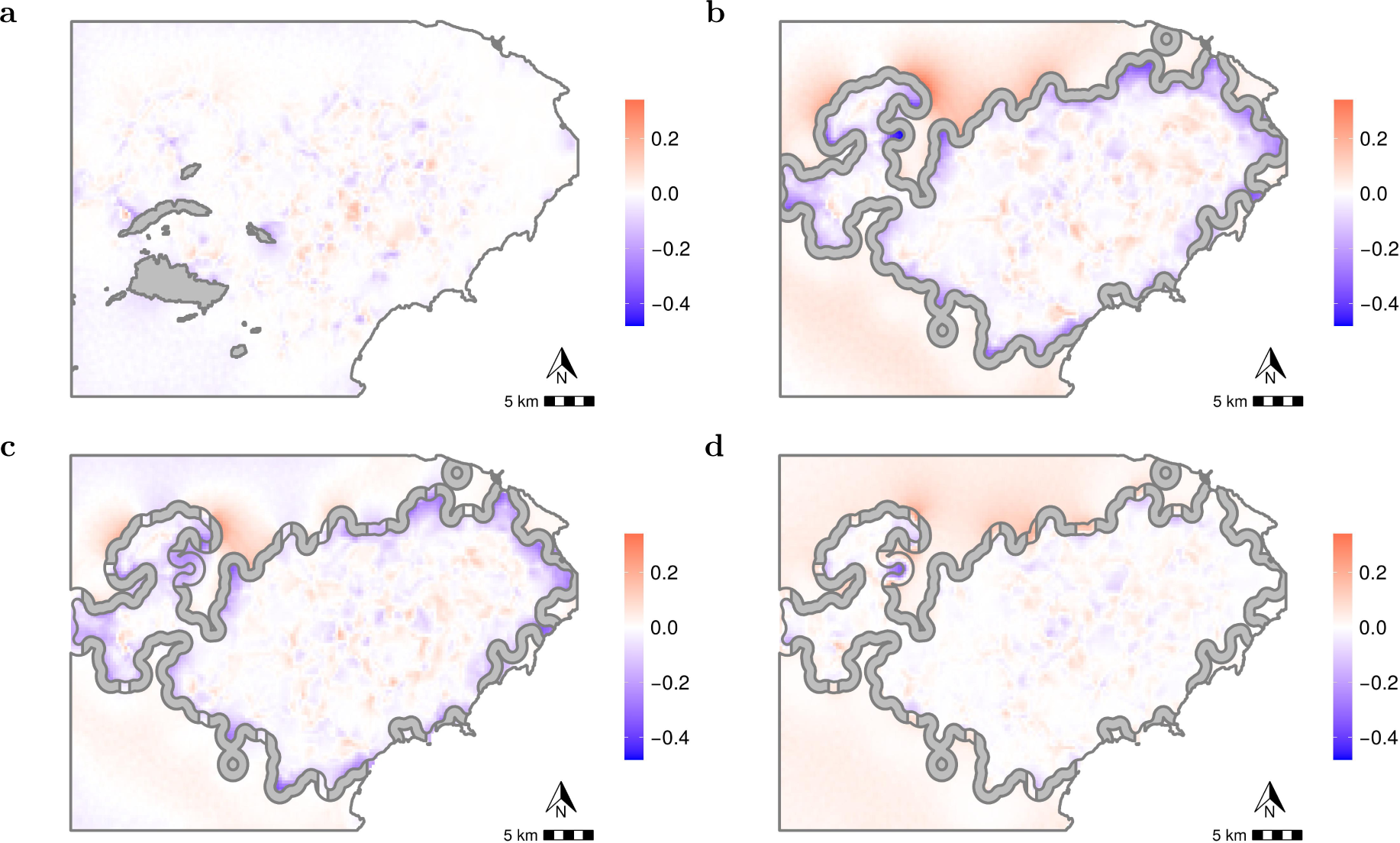
Differences in the mean of the posterior predictive distribution of the probability of *Xylella fastidiosa* presence. (a) Difference between stationary model and mountain barrier model; (b) difference between stationary model and continuous barrier model; (c) difference between stationary model and discontinuous barrier model; and (d) difference between discontinuous barrier model and continuous barrier model.

In order to help in the interpretation of the comparison of the results of the stationary model, the continuous barrier model and the discontinuous barrier model, from now on we denominate the areas on both sides of the perimeter barrier built around the positives as external and internal zones. The maximum probability of the presence of *X. fastidiosa* was 0.46 in the area corresponding to the external zone in the stationary model (Fig. 3a), while it was 0.29 and 0.36 in the continuous and discontinuous barrier models, respectively (Fig. 3e and 3g).

Considering the difference between the mean of the posterior predictive distribution of the stationary model and the continuous barrier model, only positive values were obtained in the external area of the barrier (Fig. 4b). That is, the probability of *X. fastidiosa* presence was higher with the stationary model, particularly in the northern area adjacent to the barrier. This same behavior was also observed, but to a lesser extent, when the stationary and discontinuous barrier models were compared (Fig. 4c). However, the difference between the discontinuous and continuous barrier models showed that in the areas where breaks were implemented, the probability of *X. fastidiosa* presence was similar or even increased, depending on the location. In particular, the probability of presence in the external area of the barrier increased through the breaks located in the north, while no differences were observed in those in the south-west (Fig. 4d).

With respect to the mountain barrier model, the continuous and discontinuous barrier models showed a higher probability of *X. fastidiosa* presence in the areas adjacent to the barrier (Fig. 3e and 3g). This increase in the probability of the presence of the pathogen in the internal area adjacent to the perimeter barriers was also observed in the difference between the mean of the posterior predictive distribution of the stationary model and the continuous and discontinuous barrier models (Fig. 4b and 4c).

## Discussion

The occurrence of *X. fastidiosa* in the demarcated area in Alicante was modeled using hierarchical Bayesian spatial models with the incorporation of barriers, following the methodology described by Bakka et al. (2019). Here, the main objective was to evaluate the influence of different types of barriers in the distribution of the pathogen. From the perspective of the SDMs, the presence of elements in the landscape that prevent or hinder the spread of the organisms cannot be ignored, since assuming stationarity and isotropy in this context would give inaccurate results (Bakka et al. 2019). Non-stationary models that incorporate barriers may also allow the effect of disease control interventions to be simulated.

In this case, climatic variables were not included in the models for the occurrence of *X. fastidiosa* in the demarcated area in Alicante. Previous works indicated that these variables were not relevant in this specific scenario, whereas a strong dominating effect of the spatial component was observed (Cendoya et al. 2020). Our analysis confirmed the strong spatial aggregation of *X. fastidiosa* in the demarcated area in Alicante, so the probability of *X. fastidiosa* presence was increased in the areas with higher prevalence of the pathogen compared to those where it was not detected (Fig. 3). These results are in line with other studies highlighting the importance of incorporating the spatial structure in SDMs for plant pathogens (Meentemeyer et al. 2008).

In contrast to previous studies on marine species (Bakka et al. 2019; Martínez-Minaya et al. 2019), in the case of *X. fastidiosa* the overall values of the posterior predictive distribution of the stationary model were relatively similar to those obtained with the models that incorporated barriers (Fig. 3). On the one hand, this was a somewhat unexpected result, considering that the simulated barriers were assumed to be completely impervious to the pathogen. On the other hand, the results obtained here somehow illustrate the actual difficulties involved in effectively containing the spread of the pathogen by implementing dispersal barriers (Kottelenberg et al. 2021).

Nevertheless, relevant differences in the posterior predictive distribution of the probability of *X. fastidiosa* presence resulting from the incorporation of the barriers in the models can be appreciated in finer spatial detail. When the area above an altitude of 1,065 m was considered as a barrier for the spread of *X. fastidiosa*, the main difference with respect to the stationary model was found in the zone adjacent to the barrier. In this area, the probability of the presence of *X. fastidiosa* was higher in the mountain barrier model than in the stationary model due to the smoothing effect that occurred when mountains were not considered as barriers (Fig. 4a).

The dimensions and characteristics of the cordon sanitaire, i.e. continuous or discontinuous perimeter barriers, were based on the spatial range of approximately 4 km obtained in the stationary model (Table 1). To observe differences when incorporating the perimeter barrier, it should be situated less than 4 km away from the positive samples. Due to the assumed impermeability, the width of the perimeter barriers had no influence on the probability of *X. fastidiosa* presence in the area outside the barrier. This implies that the width of the barriers used in our study cannot be interpreted in terms of the extent of the area subjected to disease control measures, such as the removal of infected host plants and vector control, as established by the Commission Implementing Regulation (EU) 2020/1201.

In the continuous barrier model, the probability of the presence of *X. fastidiosa* in the area adjacent to the outer border of the barrier was only determined by the negative samples and the intercept (Fig. 3e), resulting in a lower probability of the presence of the pathogen compared to the stationary model (Fig. 4b). Differences between the discontinuous and continuous barrier models showed that breaks in the perimeter barrier in areas with low sampling intensity, due to the greater uncertainty, resulted in a higher probability of *X. fastidiosa* presence (Fig. S3b). The increase in the probability of the presence of the pathogen through the breaks in the barrier was even greater than the difference with the stationary model (Fig. 4c). However, no major influence of the cordon sanitaire was observed in areas with a high sampling intensity adjacent to the outer border of the barrier. In those areas, the breaks in the barrier did not increase the probability of *X. fastidiosa* presence (Fig. S3c).

These results may assist plant health authorities in prioritizing the areas for the implementation of surveillance and disease control barriers. The highest priority would therefore be given to non-sampled areas close to high occurrence locations, where the implementation of a barrier would lower the probability of the presence of *X. fastidiosa*. Areas where the surveys concluded that the pathogen is absent (i.e., below the design prevalence) would be, therefore, of lower priority for the implementation of surveillance and disease control barriers, as the breaks would not increase the probability of presence of *X. fastidiosa*. These results are in line with current approaches aiming for a more targeted and risk-based management of emerging plant pathogens (Parnell et al. 2014; Hyatt-Twynam et al. 2017).

In the context of our study, the spatial aggregation obtained with the models resulted from the concurrent means of spread of *X. fastidiosa* acting during the whole time span of the epidemic. For the demarcated area in Alicante, Cornara et al. (2019) indicated that *X. fastidiosa* was detected in *P. spumarius* and *N. campestris*, with a prevalence of 27% and 1.2% of the individuals tested for the bacterium, respectively. However, the references quoted in this review do not report data on the prevalence of *X. fastidiosa* in vector populations in this region. Official samplings conducted from 2017 to 2019 by the plant health authority in the demarcated area resulted in prevalences of *X. fastidiosa* of 0.67% for *N. campestris* (n = 2,995) and 7.19% for *P. spumarius* (n = 3,157) (GVA 2020). Similar values have been reported in the Balearic Islands, with 1.12% for *N. campestris* (n = 797) and 8.25% for *P. spumarius* (n = 5,806) (MAPA 2021). However, the prevalence values in Alicante are much lower than those described for *P. spumarius* in Corsica (>40%) and Apulia (>50%) (Cruaud et al. 2018; Cornara et al. 2017; Saponari et al. 2014). These data suggest that vectors might not be playing a dominant role in the spread of the disease in the demarcated area in Alicante. Furthermore, it should be considered that the probability of infection of a plant by vectors depends not only on the prevalence, but also on the abundance of infectious vectors, their acquisition rate, transmission efficiency, the time period of the inoculation process and the infectivity of the vectors (Purcell 1981). For instance, EFSA (2019) used expert knowledge elicitation (EKE) to estimate a median acquisition rate of 12.08% and a transmission efficiency of 13.58% for spittlebug vectors in olives.

The dispersal capacity of *X. fastidiosa* vectors in Europe is rather uncertain, and no studies are available for the particular epidemiological setting in Alicante. According to a Mass-Mark-Recapture assay by Lago et al. (2020) conducted in Madrid, Spain, individuals of *N. campestris* were found at a distance of more than 2,000 m from the release point, with a relatively similar number of catches at 123 and 281 m. Studies conducted with *P. spumarius* in Apulia and Piedmont, Italy, resulted in a median dispersal from the release point of 26 m day*^−^*^1^ in an olive grove and 35 m day*^−^*^1^ in a meadow. It was estimated that 50% of the *P. spumarius* population in olives in Apulia remained within 200 m and 98% within 400 m for 2 months, with a dispersal limited to some hundreds of meters throughout the whole year (Bodino et al. 2020). EFSA (2019) conducted EKEs on the uncertainty distribution of the vector local spread and the mean distance of disease spread. The 5th, 50th and 95th percentiles of the uncertainty distribution for the vector local spread were 0.148 km, 0.767 km and 2.204 km, respectively. Percentiles for the mean distance of disease spread were 1.10 km, 5.18 km and 12.35 km, this median value being included in the 95% CI of the posterior distribution of the range of our stationary model (Table 1). This upper bound corresponds to the estimated rate of movement of the *X. fastidiosa* front in Apulia (Kottelenberg et al. 2021). Nevertheless, these EKEs were conducted under specific assumptions and their extrapolation to the scenario in Alicante is not straightforward. Among other assumptions, values were elicited for olive orchards with herbaceous cover, without the influence of competing hosts or extreme winds on vector behavior. The movement of propagating plant material was not taken into account either.

In fact, plant propagating material is considered the main pathway for the entry of *X. fastidiosa* into new regions EFSA (2019). After the introduction of the pathogen with imported infected plant material, further spread in the area can also be driven by the movement of propagating plant material. Studies reconstructing the progression of almond leaf scorch disease in Majorca indicated that *X. fastidiosa* was introduced into this island with almond buds or stems from California, and then spread through the archipelago by grafting (Moralejo et al. 2020). Grafting experiments performed in this study resulted in a transmission of about 15% with almond buds, but other studies reported values up to 60% and 80% with almond buds and stems, respectively (Mircetich et al. 1976). In the case of Alicante, genetic studies indicated that *X. fastidiosa* might also have been introduced from California (Landa et al. 2020). In the demarcated area in Alicante, almond groves were typically established with rootstock seeds that were later grafted on site with buds or stems of the scion (Cambra and Cambra 1991). These grafting materials were generally obtained from almond trees in the area or from outside when a new cultivar was first introduced. In fact, previous studies suggested that the current extent of the pathogen had arisen from a single introduction (Cendoya et al. 2020; Landa et al. 2020). Nevertheless, with the information available, it is not possible to accurately trace back the movement of propagating plant material in the area and thus determine its actual role in the spread of *X. fastidiosa*. Therefore, the spatial dependence illustrated by the models should be interpreted considering any potential means of spread, including propagating plant material and insect vectors.

The ranges obtained with the models varied from approximately 4 to 6 km (Table 1), but to relate this parameter to the actual epidemiological setting in the demarcated area in Alicante, only those from the stationary and mountain barrier models should be considered. The continuous and discontinuous barrier models incorporated simulated disease control interventions in the form of barriers, which are not present in the study area as such. Furthermore, imposing a cordon sanitaire implied a strong spatial aggregation in the area surrounded by this perimeter barrier, resulting in a greater spatial range compared to the other models studied. The models assuming no control interventions presented similar spatial dependence for the occurrence of *X. fastidiosa*. The posterior mean of the range in the stationary model was 4,030.17 m with a 95% CI (2,907.41, 5,563.88), whereas for the mountain barrier it was 3,860.88 m with a 95% CI (2,918.61, 5,212.18) (Table 1). Interpreting these values in terms of spread rates is, however, difficult as the contribution of the different means of pathogen spread cannot be disentangled. Moreover, with the information available, it is not possible to determine when the pathogen was first introduced in the area and so the temporal component is missing. Studies combining dendrochronology and phylogenetic analysis indicated that the introduction of *X. fastidiosa* in Majorca occurred around 1993 (Moralejo et al. 2020). Epidemiological models dated the introduction of the pathogen in Corsica to around 2001 when hidden infection reservoirs are not considered, and around 1985 when these non-observable hosts are included in the models (Soubeyrand et al. 2018). The rate of movement of the invasion front of *X. fastidiosa* in Apulia indicated that the disease spread started in approximately 2008 (Kottelenberg et al. 2021). Based on the low genetic diversity and the absence of recombinant events (Landa et al. 2020), it can be speculated that *X. fastidiosa* was introduced in the demarcated area in Alicante not earlier than in Majorca or Corsica.

Although spread rates cannot be inferred from our analysis, the spatial component of the models provides useful information for the management of *X. fastidiosa* in the study area. In the Matérn correlation function of the stationary and mountain barrier models, distances up to 1,792 and 1,717 m, respectively, accounted for 50% of the spatial correlation, and was less than 5% for distances longer than 5,698 and 5,459 m, respectively (Fig. 2). Regardless of the date of introduction and the weight of the different means of spread of the pathogen in the demarcated area, the mean value of the range for the stationary model indicates that host plants that were closer than 4,030.17 m to an infected plant would be at risk of giving positive for *X. fastidiosa*. Therefore, these distances should be observed to define the buffer zone where the surveillance activities will be conducted around the infested area. Originally, the Commission Implementing Decision (EU) 2015/789 established that the buffer zone surrounding the infested zone should have a width of at least 10 km. The minimum width of the buffer zone was later reduced to 5 km by the Commission Implementing Decision (EU) 2017/2352 and currently to 2.5 km by the Commission Implementing Regulation (EU) 2020/1201. Based on our models, these minimum buffer zone widths do not cover the entire area at risk for *X. fastidiosa* occurrence in the demarcated area in Alicante. Consequently, in 2019 the plant health authority implemented an additional band of 10 km surrounding the demarcated area, where official surveillance activities are also being conducted (GVA 2020).

It should be noted that the methodological improvement considering the non-stationarity of the spatial process did not increase the computational cost or the difficulty of its implementation. In fact, non-stationary models have previously been used in ecology, mainly in marine species distribution studies where terrestrial areas represent completely impervious physical barriers (Bakka et al. 2019; Martínez-Minaya et al. 2019). To our knowledge, this study is the first to apply non-stationary models with barriers in the context of plant health. However, imposing the condition that barriers are completely impermeable implies that the pathogen cannot be present or cross this area, which is a very strong assumption rarely met in practice. In the specific case of *X. fastidiosa*, these barriers represent areas without host plants and in which it is not possible for infected vectors or propagating plant material to pass through. Our discontinuous barrier model partially relaxed this assumption, allowing the pathogen to spread in some areas but still assuming that parts of the cordon sanitaire were completely impervious, which is seldom the case for *X. fastidiosa* and plant pathogens in general. Building on the present work, new modeling methods need to be developed to accommodate the incorporation of barriers with different levels of permeability, and thus more realistic plant health scenarios may be considered.

## Acknowledgements

The present work has received funding from European Union’s Horizon 2020 research and innovation programme under grant agreement no. 727987 (XF-ACTORS, “Xylella Fastidiosa Active Containment Through a Multidisciplinary-Oriented Research Strategy”), and grant E-RTA 2017-00004-C06-01 FEDER INIA AEI-MCIN and Organización Interprofesional del Aceite de Oliva Español. DC is also grateful for grant PID2019-106341GB-I00 FEDER AEI-MCIN. MC holds an IVIA grant partially funded by the European Social Fund. We thank the Plant Health Service of Generalitat Valenciana for providing the survey data. Thanks are also due to A. López-Quílez (UV), V. Dalmau and A. Ferrer (GVA) for their comments on the manuscript.

## Supporting information

Additional supporting information may be found at: Appendix_S1.pdf

## Data availability

Data and code are available at https://doi.org/10.5281/zenodo.4656029

## Notes

https://doi.org/10.5281/zenodo.4656029

### Competing Interest Statement

The authors have declared no competing interest.

