## Appendix S1 for "Barrier effects on the spatial distribution of *Xylella fastidiosa* in Alicante, Spain"

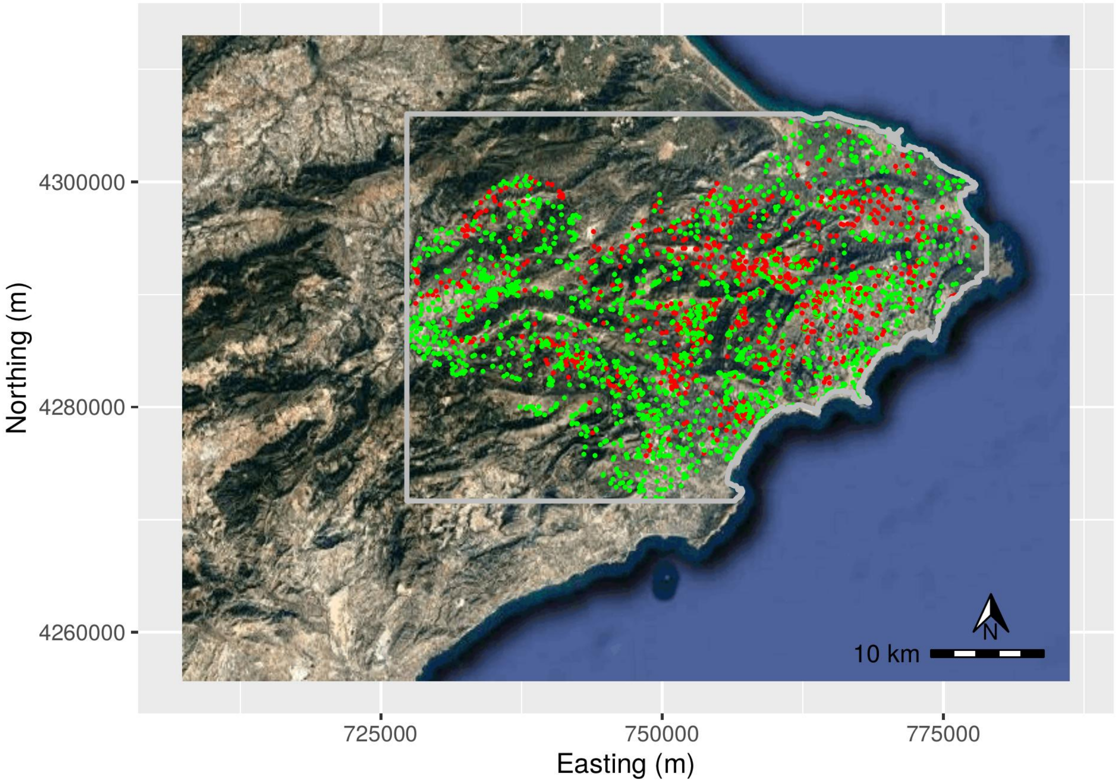


**Figure S1:** Study area (delimited by the grey line) in Alicante (Spain), in UTM coordinate system, with the positive (•) and negative (•) samples for *Xylella fastidiosa*.

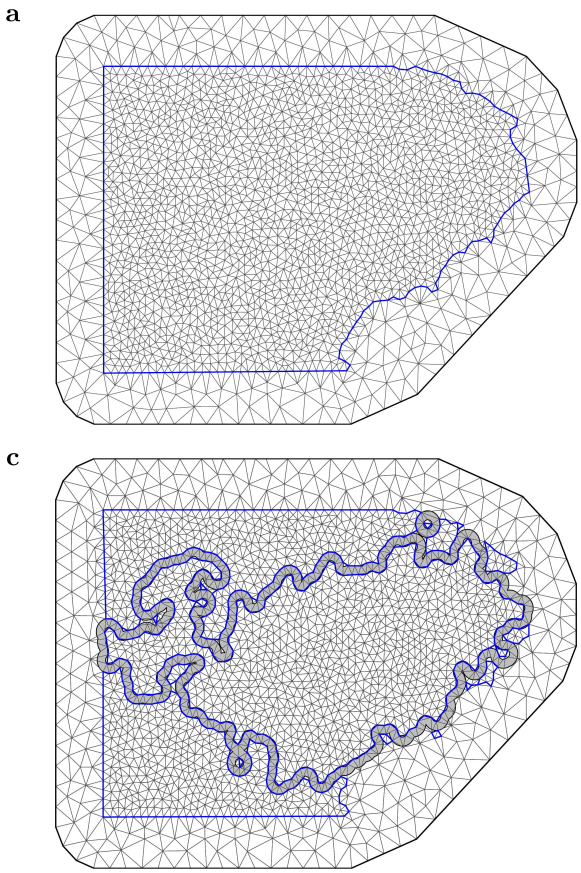


**Figure S2:** Triangulation of the study region (*mesh*) for each model. (a) Stationary model; (b) mountain barrier model; (c) continuous barrier model; and (d) discontinuous barrier model.

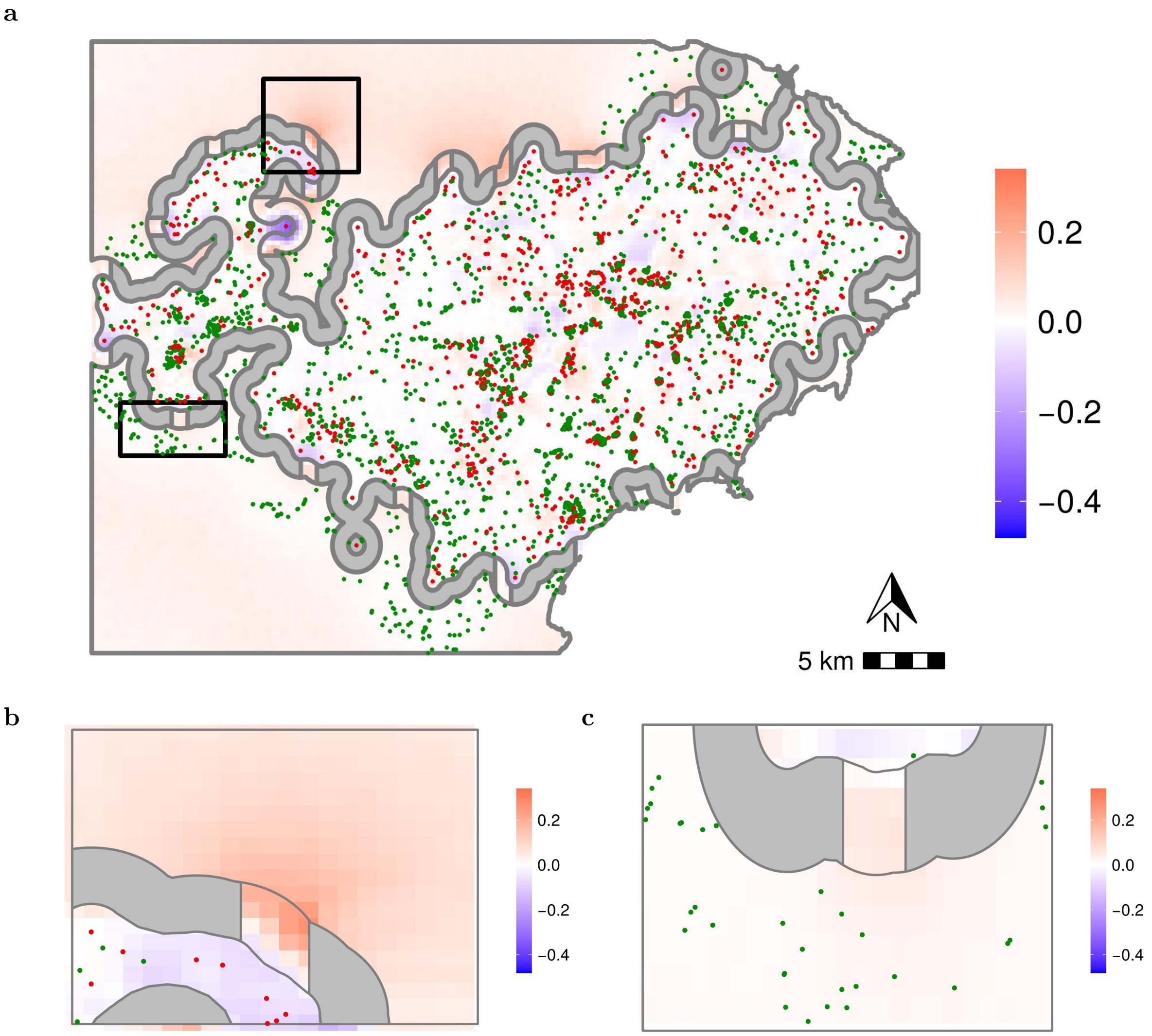


**Figure S3:** (a) Difference in the mean of the posterior predictive distribution of the probability of *Xylella fastidiosa* presence between discontinuous and continuous barrier models, with the positive (●) and negative (●) samples for *X. fastidiosa*. (b) Detail of a break with non-sampled area adjacent to the outer border of the barrier. (c) Detail of a break with high sampling intensity adjacent to the outer border of the barrier.
